# A linguistics-based algorithm for RBP motif and context discovery

**DOI:** 10.64898/2026.03.17.712526

**Authors:** Shaimae I. Elhajjajy, Zhiping Weng

## Abstract

RNA-binding proteins (RBPs) regulate their RNA targets by binding to short sequence motifs, but the underlying mechanisms enabling sequence-specific recognition within the vast transcriptome remain unclear for the majority of human RBPs. Sequence contexts are believed to be a significant contributing factor to RBP binding specificity but are often overlooked. Further, existing motif discovery algorithms do not consider the structure and composition of the motif’s flanking regions in their construction, which represents a consequential shortcoming. Herein, we present a novel linguistics-inspired RBP motif and context discovery algorithm that is consensus-based, deterministic, and flexible. Our algorithm draws multiple parallels between natural language and genomic language and relies on three important k-mer properties that impart lexical, syntactic, and semantic structures and rules to the process of motif and context discovery. Critically, our algorithm integrates information from sequence contexts when constructing RBP motifs. We demonstrate that our algorithm achieves strong discovery accuracy against a ground-truth set, and even outperforms existing methods in primary motif ranking.

## 1 Introduction

RNA-binding proteins (RBPs) are essential regulators of RNA processing, which in turn ensures proper cellular function. RBPs with structurally diverse, sequence-specific RNA-binding domains (RBDs) have been shown to recognize similar motifs, but they still demonstrate a remarkable degree of transcriptomic specificity, with each binding a distinct set of targets [1]. Because RBP motifs are relatively short in length (3-8 nucleotides [nt] [2]) and low in nucleotide complexity [1], binding sites generally lack substantial variety, suggesting that other sequence-based and biological features facilitate the discriminative binding ability of RBPs. While previous studies implicate sequence context as a contributing factor [1], their precise role remains unclear, and their involvement has not been established at large scale *in vivo*.

Motif discovery is a challenging problem due to complexities introduced by a vast genomic search space and RBP motif degeneracy [3] [4] [5] [6]. Conventional motif discovery algorithms, including statistical and probabilistic methods, suffer from a lack of differentiation between over-and under-represented motif instances, as well as from their disuse of structural features and relationships between sequence components. These limitations allow the admission of noise, such as insignificant *k*-mers, into the final motif, results in either poor approximation or erroneous motifs altogether.

In our previous work, we formulated a novel linguistics-based representation and modeling of RBP binding prediction that identifies strongly salient regions with high accuracy [7]. Continuing our work, we herein present a novel motif discovery algorithm that enables accurate, deterministic, and fast identification of RBP motifs and contexts. In addition to being consensus-based, our approach is context-aware, leveraging the properties and relationships among *k*-mers, their associated regions, and their sequences of origin. We demonstrate that our algorithm discovers RBP motifs with very strong accuracy. Importantly, our algorithm enables us to discover novel *in vivo* sequence contexts and nucleotide preferences of RBP binding for 71 RBPs in HepG2 and 74 in K562, with consistent results across cell lines denoting the robustness of our method. Taken together, our work facilitates the investigation of contextual determinants of RBP binding specificity, which will help to improve our understanding of RNA regulation.

## 2 Methods

### 2.1 Objectives, Requirements, and Assumptions

To address the limitations of conventional algorithms and to pursue our aims of accurate RBP motif and context discovery, we built upon our previous work to develop a novel linguistic-based motif discovery algorithm. To this end, we set forth a series of objectives that guided our formulation and design, requiring that our algorithm satisfy the following predefined requirements: (1) Flexibility, to accommodate our contexts data structure, discover RBP binding motifs and contexts directly from our MIL predictions, and achieve motif and context discovery at different levels of granularity and generalizability through simple changes in configuration and parameter values; (2) Deterministic, without integration of stochastic techniques, to ensure that our discovered motifs and contexts are stable and consistent for each run; (3) Consensus-based, to vastly reduce the motif search space during discovery, and to avoid errors caused by the assumption that *k*-mer overrepresentation alone warrants its membership to a motif; (4) Context-aware, to leverage the composition and properties of the flanking regions, as well as similarity and co-occurrence relationships between motif instances and context units; (5) Unbiased, to treat every enriched *k*-mer as a candidate motif consensus, thus enabling the discovery of all possible motifs within a dataset by relying only on the structure, composition, and biological context of the sequences in the target RBP’s dataset; (6) Accurate, to ensure high-confidence discovery of known and novel motifs, especially for RBPs with little characterization, and to ensure correct context discovery by preventing noise propagation; and (7) Efficient, to improve time complexity and enable fast motif discovery with a parallelizable implementation in mind.

The conceptualization, formulation, and implementation of our algorithm were guided by the following axioms and assumptions: (1) Our dataset consists of a foreground of positive sequences and a background of negative sequences; (2) Enriched *k*-mers occur in greater abundance in positive sequences with respect to negative sequences; (3) Enriched *k*-mers occur in clusters to form a region significant to binding; (4) Positive sequences may contain more than one significant region; (5) Each positive sequence contains only one putative motif instance; (6) An RBP binding site is comprised of enriched *k*-mers; (7) *k*-mers comprising a binding motif or context are enriched; (8) An RBP binding motif has an associated motif consensus; (9) An RBP binding motif is composed of a set of motif instances; (10) Putative motif instances are significantly enriched in positive sequences with respect to negative sequences; (11) Significant *k*-mer enrichment is not a sufficient condition to qualify as a motif instance; (12) Putative motif instances possess some degree of sequence conservation with both the motif consensus and with each other; (13) Putative motif instances share some degree of co-occurrence with the motif consensus; (14) Putative motif instances have significantly enriched contexts.

### 2.2 A natural language- and linguistics-inspired representation of RBP motifs and contexts

The overarching goals of our work are to provide RNA sequences and RBP binding sites with a linguistic representation, and to leverage principles from natural language to emulate techniques of linguistic analysis to discover, analyze, and understand RBP binding patterns (**Figure 1**). Accordingly, in our previous work, we have defined lexical, syntactic, and semantic constructs to model RBP-bound RNA sequences [7]. In brief, at the lexical level, we consider *k*-mers to be the fundamental lexical units, analogous to words in natural language (**Figure 1A**, left), and we consider these *k*-mers to have hierarchical lexical meaning: at the first level, discerning between enriched and non-enriched *k*-mers; at the second level, determining whether enriched *k*-mers have roles as motif or contextual units; and at the third level, determining whether each motif unit is a motif consensus or motif instance (**Figure 1B**). At the syntactic level, we map regions to phrases and sequences to sentences. We then define a syntactic form, consisting of a central target *k*-mer situated between two flanking regions, that introduces syntactic structure to regions and sequences (**Figure 1C**), and we identify the most significant regions within a sequence that contribute to the resulting RBP binding event (**Figure 1A**, middle). At the semantic level, we consider *k*-mer overrepresentation and region significance as proxies for lexical and phrasal semantics, respectively.

**Fig. 1.**
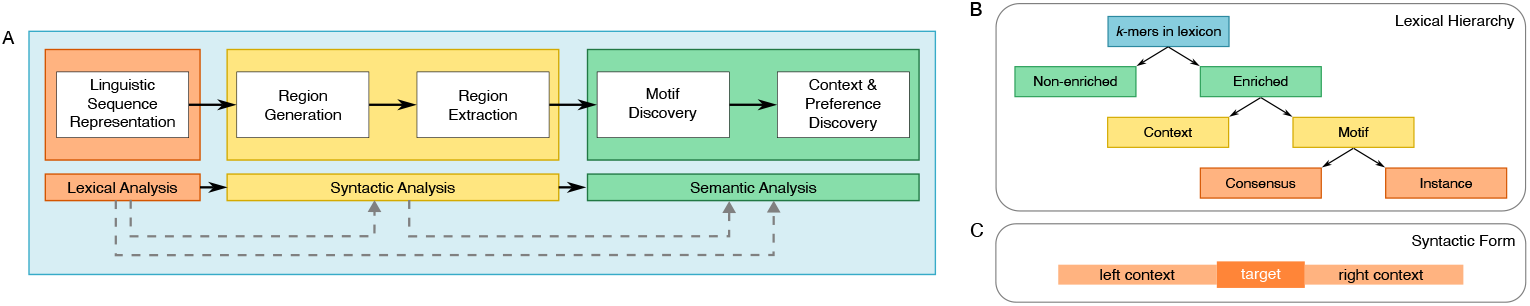
Linguistic-inspired workflow, stratified by level of linguistic analysis. (A) Our linguistic representation and formulation of our defined lexical and syntactic analysis, as in [7]. Our work herein covers semantic analysis, as represented by motif and context discovery. (B) Hierarchical lexical semantics. (C) Syntactic form, consisting of a central target *k*-mer and its flanking regions.

In this work, we primarily focus on the semantic aspects of our approach while leveraging previously defined lexical and syntactic representations (**Figure 1A**, right). We emulate additional linguistic concepts to model structures and relationships in RBP-bound RNA sequences by defining semantic rules to analyze sought-after motifs and contexts. First, we define *k-mer enrichment* to be the measure of *k*-mer overrepresentation in bound sequences relative to unbound sequences, and we consider this enrichment to approximate lexical frequency, a marker of word importance. Second, we define *k-mer similarity* as analogous to word synonymy, which enables the analysis of semantic relationships between *k*-mers; we consider *k*-mers that are members of a motif as synonyms. Third, we define *k-mer co-occurrence* as the degree to which two *k*-mers occur together within the same sentence; this property serves as a parallel for lexical co-occurrence, which is used as a proxy for semantic relationships, where frequently co-occurring words are considered to be strongly associated [8, 9].

### 2.3 Problem definition and formulation

We formulate the RBP motif discovery problem as follows: given a set of sequences and a candidate motif consensus, find the set of *k*-mers within the sequences that fulfill the following conditions:

1. Each *k*-mer is significantly enriched.
2. Each *k*-mer has sufficient sequence similarity with the candidate motif consensus.
3. Each *k*-mer satisfies a co-occurrence requirement with the candidate motif consensus.
4. When all *k*-mers in the set are aligned, the consensus of the resulting motif is equivalent to the starting candidate consensus.

We developed a novel motif discovery algorithm that leverages three important *k*-mer properties – *k*-mer enrichment, *k*-mer similarity, and *k*-mer co-occurrence – to derive RBP binding motifs. The algorithm is composed of 6 stages, **Algorithm 1**).

#### Definition 1

Given a set of labeled sequences *X* = {(*X*_1_, *Y*_1_), (*X*_2_, *Y*_2_), …, (*X*_*n*_, *Y*_*n*_)}, we decompose each sequence *X*_*i*_ [1 ≤ *i* ≤ *n*] into *m* regions called “contexts”, *C*_*i*_ = {*C*_*i*1_, *C*_*i*2_, …, *C*_*im*_} [7]. Each context *C*_*ij*_ [1 ≤ *j* ≤ *m*] has an associated label *Z*_*ij*_ = *Y*_*i*_, initialized by label inheritance.

#### Definition 2

*f*_*C*_ is a binary contexts classifier trained with iterative relabeling [7], producing predicted labels 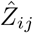 for each *C*_*ij*_ to approximate the true labels *Z*_*ij*_, where:

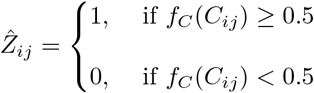

Let 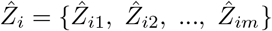 be the instance-level prediction labels corresponding to contexts *C*_*i*_ [1 ≤ *i* ≤ *n*]. The full set of contexts and their predictions across all sequences are denoted 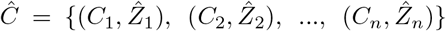, where 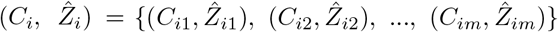.

#### Definition 3

We denote 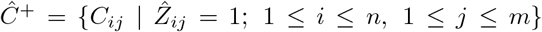 and 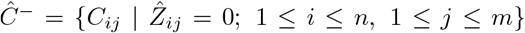 as the sets of contexts predicted as positive and negative, respectively, by *f*_*C*_, and *Ĉ* = *Ĉ*^+^ ∪ *Ĉ*^−^ denotes the union of both types of contexts.

### 2.4 A novel motif discovery algorithm to discover RBP motifs and contexts

Our algorithm design decisions rely on the use of context predictions as a method of extracting significant regions, together with the composition and structural properties of related sequences, to discover motifs using a consensus-based approach that requires three important empirically-derived *k*-mer properties supported by theoretical, statistical, probabilistic, linguistic, and biological foundations. The algorithm (1) utilizes ***k* -mer enrichment** to reduce the motif search space by filtering out insignificant *k*-mers that are unlikely to be candidate motifs (**Algorithm 1**, lines 1-2), (2) uses ***k* -mer similarity** to further reduce the motif search space by filtering out motif instances with inadequate sequence conservation properties with respect to a candidate consensus (**Algorithm 1**, line 5), and (3) employs ***k*-mer co-occurrence** constraints to retain only the most probable motif instances for each candidate consensus (**Algorithm 1**, line 6). Importantly, these 3 *k*-mer properties are each necessary but not independently sufficient for accurate motif discovery; only in conjunction are they necessary and sufficient conditions. The motif itself is constructed at the end of the algorithm by identifying all occurrences of discovered motif instances along with their respective contexts (**Algorithm 1**, lines 9-10), and the optimal motif is selected based on an iterative scoring strategy (**Algorithm 1**, line 12). The algorithm consists of 6 stages, as follows (**Figure 2**).

**Fig. 2.**
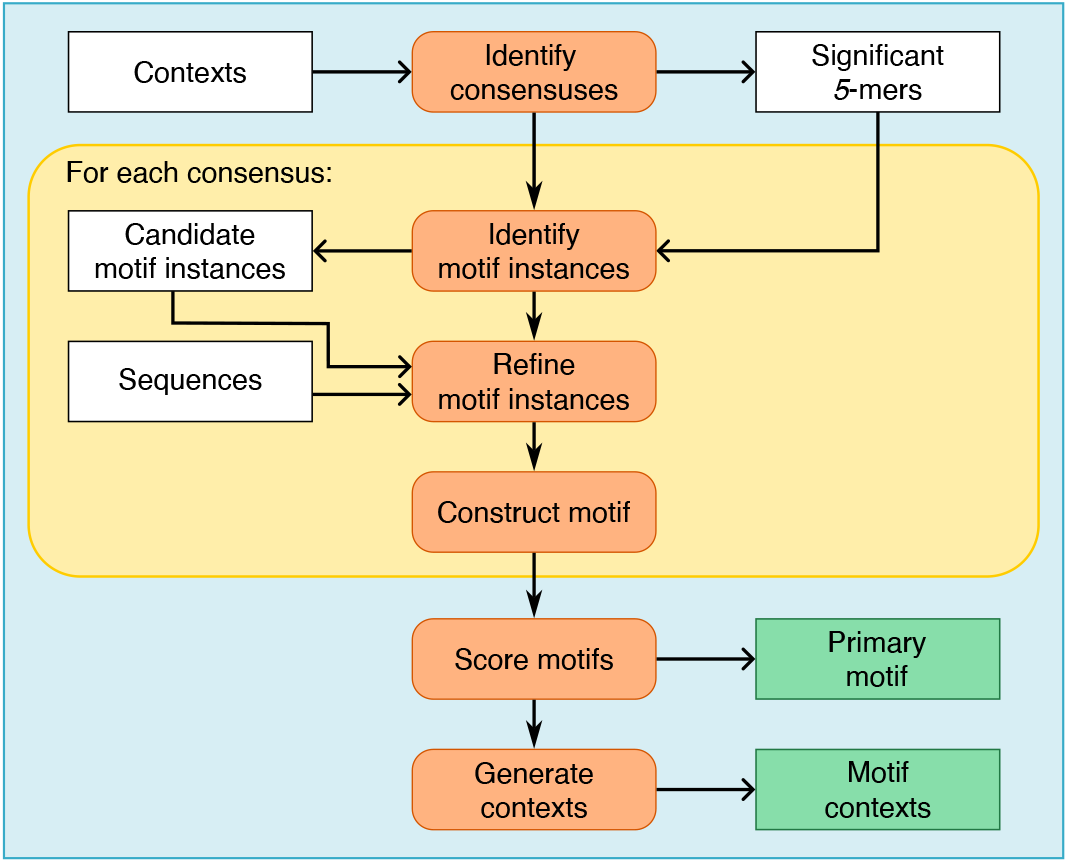
Process diagram representing the conceptualization of our six-stage motif discovery algorithm. The operations contained within the yellow box indicate those which are performed for each candidate motif consensus independently.

#### 2.4.1 Stage I: Identification of candidate motif consensuses

In the first stage, we sought to identify the most highly probable consensus *k*-mers of candidate motifs. We reasoned that a candidate motif consensus would have two main attributes: (1) being the target of a context with a high predicted class probability, which suggests that it contains salient features highly significant to RBP binding, and (2) being strongly enriched, which indicates that these *k*-mers are overrepresented in RBP binding sites, in agreement with our initial assumptions. As such, let Σ = {*A, C, G, U*} be the RNA alphabet, and let *L* = {*l*_1_, *l*_2_, …, *l*_*s*_} represent the set of all possible *k*-mers, such that *s* = |*L*| = |Σ|^*k*^. We consider L to contain the initial set of candidate motif consensuses, and we set *k* = 5, which has previously been established as a representative *k*-mer size for RBP binding [1]; thus, our dictionary size is *s* = 4^5^ = 1024.

##### Identification of local maxima

For each sequence *X*_*i*_, we considered the predictions of its constituent contexts 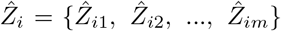 as a continuous function, then identified all local maxima *θ*_**i**_ across that function together with their corresponding targets. Generalizing across all sequences, we compile all such target *k*-mers:

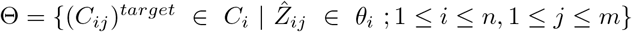

##### Computation of *k*-mer enrichments

For each *k*-mer *l*_*q*_ in *L* [1 ≤ *q* ≤ *s*], let 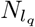 and 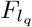 represent the count and frequency, respectively, of *l*_*q*_’s occurrence as a target in a given set of contexts. We computed its corresponding enrichment 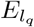 by calculating the frequency of its occurrence as a target in positively-predicted contexts *Ĉ*^+^ normalized by the frequency of its occurrence as a target in negatively-predicted contexts *Ĉ*^−^, as follows:

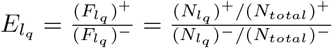

where (*N*_*total*_)^+^ = |*Ĉ*^+^| and (*N*_*total*_)^−^ = |*Ĉ*^−^|. We then performed chi-square testing to obtain a *p*-value for each target *l*_*q*_ that represents the statistical significance of its enrichment 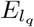.

##### Filtering candidate motif consensuses

We filtered *L* to only retain the most probable candidate motif consensuses by applying two conditions:

*Condition 1*: A candidate motif consensus must be the target of a context corresponding to a local maximum. In other words, we retain only those *k*-mers in *L* that are also in Θ.

*Condition 2*: A candidate motif consensus must be enriched. In other words, we retain only those *k*-mers in *L* for which 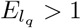.

Let Λ denote the set of *k*-mers that satisfies both of the above conditions. Then:

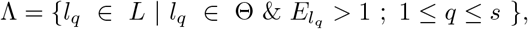

where we rename the elements of Λ as {*λ*_1_, *λ*_2_, …, *λ*_*t*_}, such that *t* ≤ *s*; in other words, Λ ⊂ *L*. Λ represents the final set of candidate motif consensuses to be considered for motif discovery.

#### 2.4.2 Stage II: Similarity partition construction to determine candidate motif instances

In the second stage, we aimed to account for both the relative sequence similarity observed between a motif’s consensus and related instances [5, 6] and the degenerate nature of RBP motifs [10]. For each candidate consensus *λ*_*r*_ in Λ [1 ≤ *r* ≤ *t*], we assembled a preliminary list termed a “partition”, *P*_*r*_, containing all enriched *k*-mers with restricted sequence similarity to that consensus; that is:

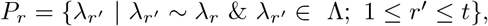

where the symbol ∼ indicates sufficient sequence similarity as defined in what follows.

We constructed this preliminary partition on the basis of similarity to the corresponding consensus. We first generated 5 partitions, *P*_1_ through *P*_5_, each containing all other 5-mers in Λ that share the same nucleotide as candidate consensus *λ*_*r*_ at the same position, where indices 1 through 5 denoting the partition number correspond to the position of concern. We then merged these lists in a step-wise fashion: first, by intersecting the lists corresponding to the inner positions (*P*_*inner*_ = *P*_2_ ∩ *P*_3_ ∩ *P*_4_), resulting in a maximum of 4^2^ = 16 possible 5-mers; second, by intersecting the lists corresponding to the outer positions (*P*_*outer*_ = *P*_1_ ∩ *P*_5_), resulting in a maximum of 4^3^ = 64 possible 5-mers; and third, by performing the union on the outcomes of the two intersections (*P*_*r*_ = *P*_*inner*_ ∪ *P*_*outer*_), resulting in a total maximum of 16+64 = 80 possible 5-mers. This yielded a preliminary partition, *P*_*r*_, consisting of *k*-mers with at most 3 substitutions with respect to the candidate consensus *λ*_*r*_, thus modeling the degeneracy of RBP motifs. Note, however, that the partition does not contain all possible *k*-mers with at most 3 substitutions with respect to its consensus. *P*_*r*_ served as the initial partition for candidate consensus *λ*_*r*_ – pending further filtering – containing the corresponding motif’s potential instances with sufficient sequence similarity to the candidate consensus.

##### A remark on modeling motif conservation

A conventional formulation of motif discovery is the (*k, d*)-motif search [11]: under the general assumption that motifs have some degree of sequence conservation, identify all instances of a given motif *M* implanted within a set of sequences *X*, such that the following criteria are satisfied:

1. The motif M is of length *k*, has the consensus *µ*_0_, and is composed of a set of instances {*µ*_1_, *µ*_2_, …, *µ*_*H*_}.
2. Each sequence *X*_*i*_ [1 ≤ *i* ≤ *n*] in *X* contains exactly one instance of the motif.
3. Each instance *µ*_*h*_ [1 ≤ *h* ≤ *H*] differs from the consensus *µ*_0_ by no more than *d* nucleotides.

A variant of the (*k, d*)-motif search introduces a quorum constraint, which alternatively requires only a subset of sequences in X to contain a motif instance [12].

The Hamming Distance (HD) is used to measure the position-specific sequence dissimilarity (i.e., the number of mismatches) between any two strings [13]:

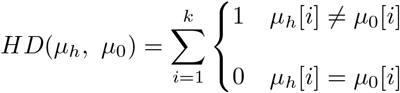

The set of all *k*-mers satisfying criterion #3 above is called the “*d*-neighborhood” of consensus *µ*_0_, which can be represented by *V*_*d*_(*µ*_0_) = {*µ*_*h*_ | *HD*(*µ*_*h*_, *µ*_0_) ≤ *d* ; 1 ≤ *h* ≤ *H*} and whose size can be modeled by [11]:

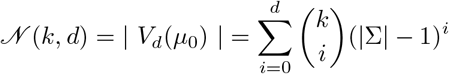

Let *k* = 5 and *d* = 3. Then the 3-neighborhood for a consensus of length 5 has 376 5-mers. In other words, there are 376 possible 5-mers that have at most 3 mismatches with respect to the motif consensus, though not all will be enriched. This represents a relatively wide search space from which to identify putative motif instances.

The (*k, d*)-motif search is limited by its treatment of *d* as the maximum number of tolerable mutations that a *k*-mer can possess in order to qualify as a potential motif instance for a given consensus. In reality, however, many genuine motif instances will have fewer than *d* mutations; consequently, larger values of *d* will naturally see a larger number of spurious *k*-mers satisfying the *d* condition, thus considerably expanding the motif search space [14]. Our consensus-based algorithm resembles the (*k, d*)-motif search in its consideration of sequence similarity, but introduces constraints that help reduce the search space by 4.7-fold, from 376 (using a Hamming Distance requirement in the conventional (*k, d*)-motif search) to 80 (using our approach). We demonstrate that this search space reduction is still able to efficiently produces accurate motifs.

#### 2.4.3 Stage III: Refinement of motif instances with *k*-mer co-occurrence

While the previous filtering conditions are necessary for motif construction, they are insufficient for accurate motif discovery because (1) *k*-mers may have high prediction scores and high enrichment values if they are overrepresented in the contexts of RBP binding sites, but they may not necessarily be a part of the motif itself, and (2) *k*-mers may satisfy the general requirement of sequence similarity with respect to a given candidate consensus, but the precise positions of similarity may not correspond to the most highly conserved nucleotides of the motif. RBFOX2 serves as a prime example in both regards: G-rich *k*-mers were highly enriched in the RBFOX2 dataset, but they are not constituents of the RBFOX2 motif due to their role in RBFOX2 binding contexts, which are known to be G-rich [1]; and *k*-mers may exhibit sufficient overall sequence similarity to RBFOX2’s canonical GCAUG motif, but may not share the essential G1, U4, and G5 positions, which are largely immutable for RBFOX2 binding [15]. We thus required an additional constraint to identify only the most highly-probable instances for a given motif.

##### Introducing *k*-mer co-occurrence

Because each RBP binding site represents a single binding event, we hypothesized that multiple instances of the RBP’s motif would not frequently occur within the same sequence. Taking into consideration that motif instances may cluster together, our assumption dictates that only one binding event occurs at a time for a particular RBP at a site at which a particular motif instance is located. This hypothesis in turn informed our assumption that there may be some as-of-yet undefined co-occurrence relationship between a motif consensus and its motif instances. We observed in our data an intriguing relationship between a motif consensus and any putative motif instance, whereby the two *k*-mers occurred within the same 101 nt long sequence at some frequency. We termed this feature the proximal *k*-mer co-occurrence, which carries a significant property that defines a *k*-mer as a true motif instance. We defined the *k*-mer co-occurrence as the frequency at which a particular *k*-mer *ρ*_*a*_ [1 ≤ *a* ≤ *u*] (representing a potential motif instance) in a given partition *P*_*r*_ occurs in the same sequence as its corresponding candidate consensus *λ*_*r*_:

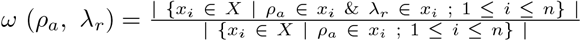

In keeping with our linguistic analogies, the co-occurrence between *k*-mers is similar in principle to the linguistic concept of lexical co-occurrence, which uses the degree to which words co-occur within the same sentence as a proxy for word association and semantic relationships [8, 9]. Similarly, we posit that the degree of co-occurrence observed between a motif instance and its consensus carries some biological, functional connotations for further exploration.

##### Filtering partitions using *k*-mer co-occurrence

In the third stage, we filtered partitions of *k*-mers that were less likely to be true motif instances by removing any *k*-mers whose co-occurrence frequency was above a pre-defined co-occurrence threshold and, consequently, retaining only those *k*-mers that satisfied this threshold. To determine a suitable value for this threshold in an unbiased manner, we devised a minimization-based tuning algorithm.

Formally, for a given candidate consensus *λ*_*r*_, we computed the co-occurrence *ω*(*ρ*_*a*_, *λ*_*r*_) for each *k*-mer *ρ*_*a*_ in its initial partition *P*_*r*_. We only retained *k*-mers in *P*_*r*_ with co-occurrence frequency *ω*(*ρ*_*a*_, *λ*_*r*_) *> ϕ*, where *ϕ* represents a designated co-occurrence threshold. We tested multiple values of *ϕ* and determined the optimal one by tuning until convergence using the following procedure.

We initialized the tuning algorithm by filtering *P*_*r*_ based on a starting co-occurrence threshold of *ϕ* = 0.5, then comprehensively tested values of *ϕ* in both numerical directions (**Algorithm 2**, lines 1-7). We first tuned in the upwards direction by iteratively increasing *ϕ* by a small increment *ϵ* until a ceiling at 1. At each incremented value of *ϕ*, we performed the following operations:

1. Generated a filtered partition Π_*r*_ by removing any *k*-mers from *P*_*r*_ whose co-occurrence did not satisfy the threshold (i.e., for which *ω* (*ρ*_*a*_, *λ*_*r*_) *> ϕ*) (**Algorithm 3**, line 5)
2. Constructed the candidate motif based on the motif instances in Π_*r*_ (**Algorithm 3**, lines 6-7) (**Section 2.4.4**)
3. Computed the Kullback-Leibler Divergence (KLD) between the position probability matrices (PPMs) of candidate motifs from two successive iterations of tuning (**Algorithm 3**, line 8)
4. Decided whether to (a) stop tuning, if the successive KLDs began to increase, or (b) continue tuning, if the successive KLDs continued to decrease (**Algorithm 3**, lines 9-15)

In step 3, we treat motif PPMs as a sequence of probability distributions and use the KLD to determine how similar the PPMs from two successive iterations are. Let *P* represent the PPM from some tuning iteration *i*, and let *Q* represent the PPM from the next tuning iteration *i* + 1; then, the KLD is computed by:

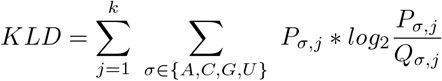

By continuing to tune the co-occurrence threshold as long as successive KLDs continued to decrease, we minimized the difference between probability distributions of motifs at successive co-occurrence thresholds. In this manner, we treat the series of related PPMs as a set of points that cluster together around a special point called a limit point, and the final PPM will be a very close approximation of the limit point, if not in fact the actual limit point.

We repeated this tuning procedure in the downwards direction by iteratively decreasing *ϕ* by a small decrement −*ϵ* until a floor of 0. To arrive at the final co-occurrence threshold, we selected the value of *ϕ* associated with the smallest difference in KLD between the two tuning directions. (**Algorithm 2**, lines 8-12). The resulting partition after co-occurence filtering, Π_*r*_, served as the final partition containing all unique motif instances {*π*_1_, *π*_2_, …, *π*_*v*_} associated with the candidate motif having consensus *λ*_*r*_.

#### 2.4.4 Stage IV: Motif construction

In the fourth stage, we constructed motifs using the final partitions from Stage III. For each candidate consensus *λ*_*r*_, for each unique *k*-mer in its final partition Π_*r*_, we identified all positively-predicted contexts having that *k*-mer as a target. In accordance with our assumptions, we also implemented filtering criteria to ensure only one motif instance per sequence was used to construct the motif. If more than one instance occurred as the target of contexts originating from the same sequence, we retained only one instance using the following conditions: (1) if the identities of the instances were different, we retained the one with the higher enrichment; (2) if the identities of the instances were the same, we retained the one with the higher prediction score. We compiled and aligned all extracted instances to build the final motif *M*_*r*_ for the given consensus *λ*_*r*_.

We next computed a PPM for the motif, where the frequency for a nucleotide *σ* at a given position *j* is calculated by:

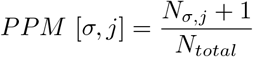

where *N*_*σ,j*_ represents the number of nucleotides at position *j* in the motif whose identity is *σ* and *N*_*total*_ represents the total number of nucleotides at position *j*; a pseudocount of 1 in the numerator prevents probabilities of zero [13].

#### 2.4.5 Stage V: Motif scoring and primary motif selection

Our algorithm discovers all possible motifs within a given RBP dataset that satisfy the algorithm’s conditions; the number of discovered motifs is different for each RBP. In the fifth stage, we select a primary motif for each RBP by using a novel, multi-metric, iterative scoring strategy based on *k*-mer and motif metrics to rank all discovered motifs.

##### A remark on weighted relative entropy

The relative entropy (RE; also referred to as the Kullback-Leibler Divergence) quantifies how well one probability distribution can approximate another; in other words, it quantifies the similarity between two probability distributions [16]. In motif discovery applications, the relative entropy measures how likely it is that a given motif, represented by a PPM for which each matrix element contains the probability *p*_*σ,j*_ of observing nucleotide *σ* at position *j* of the motif, is not produced by the background nucleotide distribution *b* [3, 5, 6]:

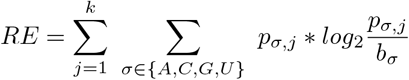

While relative entropy successfully accounts for inherent nucleotide biases within genomic sequences by correcting for the background distribution, it cannot be used to comparatively evaluate motifs that are each constructed from a different number of motif instances [3]. Hence, we instead relied on a previously-reported metric that is analogous to a log-likelihood ratio, computed by multiplying a motif’s relative entropy by its total number of constituent motif instances *w* (i.e., the weight) [3].

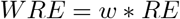

We term this the “weighted relative entropy” (WRE), with the incorporation of the weight making motif comparisons possible.

##### Primary motif selection using weighted relative entropy

For a given RBP, our approach yields every possible motif that fulfills the conditions of the algorithm. This is in fact advantageous in two ways, as it allows for the possibility of discovering (1) potential unknown secondary motifs that are bound with lower affinity by the RBP and (2) potential motifs recognized by other RBPs that may be implicated in RBP-RBP interactions. Further, because our approach is consensus-based, we discover every possible motif in a dataset, which poses a challenge when seeking to determine the primary motif for an RBP, as no single metric is fully expected to identify the correct motif [6]. We employed a multi-step scoring strategy, integrating a combination of metrics to rank the resultant motifs and select the strongest, most probable motif as the primary motif (**Figure 3, Algorithm 4**).

**Fig. 3.**
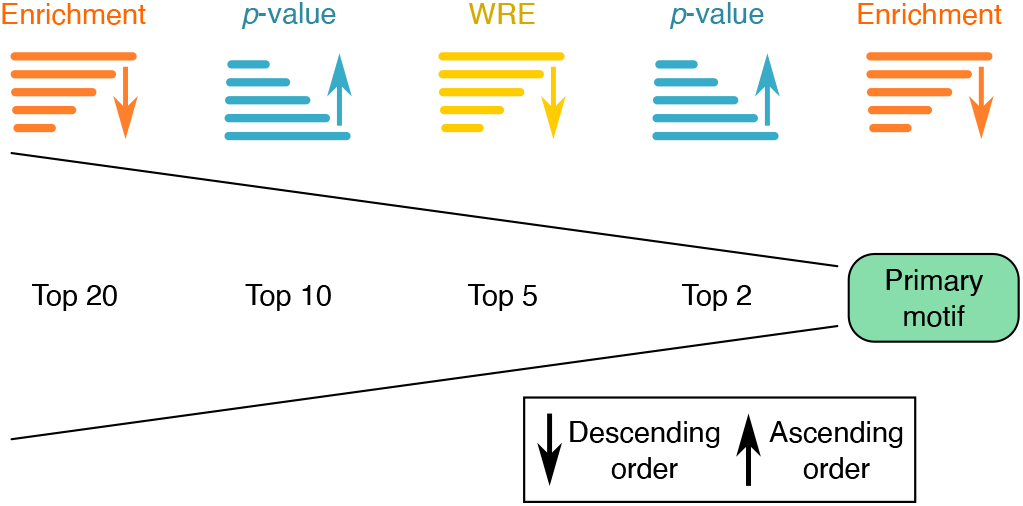
Motif scoring and primary motif selection. Motifs for each candidate consensus are ranked using a multi-metric iterative scoring strategy.

For a particular RBP, we used *k*-enrichment as the first metric to select the top 20 candidate consensuses with the highest enrichments. We then used *p*-value as the second metric to sort these candidate consensuses and selected the top 10 most significant. To narrow down the candidates further, for the 10 most significantly enriched candidate consensuses, we used WRE as the third metric, computing the WRE for each corresponding motif and then selecting the top 5 with the highest WRE as potential motifs for the RBP in question. To appoint the primary motif, we subset the top 2 of the 5 whose consensus had the smallest *p*-value, then selected from these the motif whose consensus was most highly enriched in the context predictions.

Importantly, we note that high-scoring but lower-ranking motifs still represent biologically viable RBP motifs, which we hypothesize may serve as secondary motifs for the associated RBP that may be bound, for example, at lower affinities, in specific biological contexts, or for targeted biological functions; in addition, they may be candidates for the motifs of interacting RBPs.

#### 2.4.6 Stage VI: Context discovery

In the sixth and final stage, we extracted the sequence contexts of our discovered motifs. We noticed that the position of identified motif instances within their original sequences was variable, with the vast majority occurring in the sequence’s interior and a minority occurring at or in the proximity of either end of the sequence. These boundary cases, in which a motif instance was located at either of the sequence’s extremities, made it infeasible to use the original context definitions to build the context logo, as it would result in an incomplete left or right context. To account for these boundary cases, we instead utilized the reference genome to extract the genomic coordinates of the left and right flanking regions (*±* 25-nt) surrounding each motif instance using symmetrical extension. We then aligned the resulting 55-nt long regions, computed its PPM in the same manner as previously described (**Section 2.4.4**). Note that we implemented the context width as a customizable parameter of the algorithm to enable analysis of shorter or longer sequence contexts, as desired.

### 2.5 Visualization of motifs, contexts, and binding preferences

To visualize our motifs as a sequence logo, we aligned all discovered motif instances and ran both WebLogo [17] and LogoMaker [18]. We repeated the same procedure for our discovered contexts. The resulting logos allow visualization of the degree of overall and position-specific conservation for both motifs and contexts. To further visualize the nucleotide preferences of RBP binding, we used the motif and context PPMs to generate line plots of position-specific nucleotide frequencies across the full length of the motifs and contexts.

## 3 Results

### 3.1 Motif discovery algorithm achieves strong, robust performance

Following a comprehensive literature review, we assembled a ground-truth set of 14 RBPs in each cell line that have extensively and consistently characterized motifs, then evaluated our discovered motifs relative to this ground-truth set (**Figure 4**). In HepG2, we successfully discovered 13 of 14 RBPs, leading to an accuracy of 92.86%. Our discovered motif for PABPN1 in HepG2 had a GCUGG consensus, which did not correspond with its canonical poly(A) motif [19]; we have addressed this apparent discrepancy in previous work [20].

**Fig. 4.**
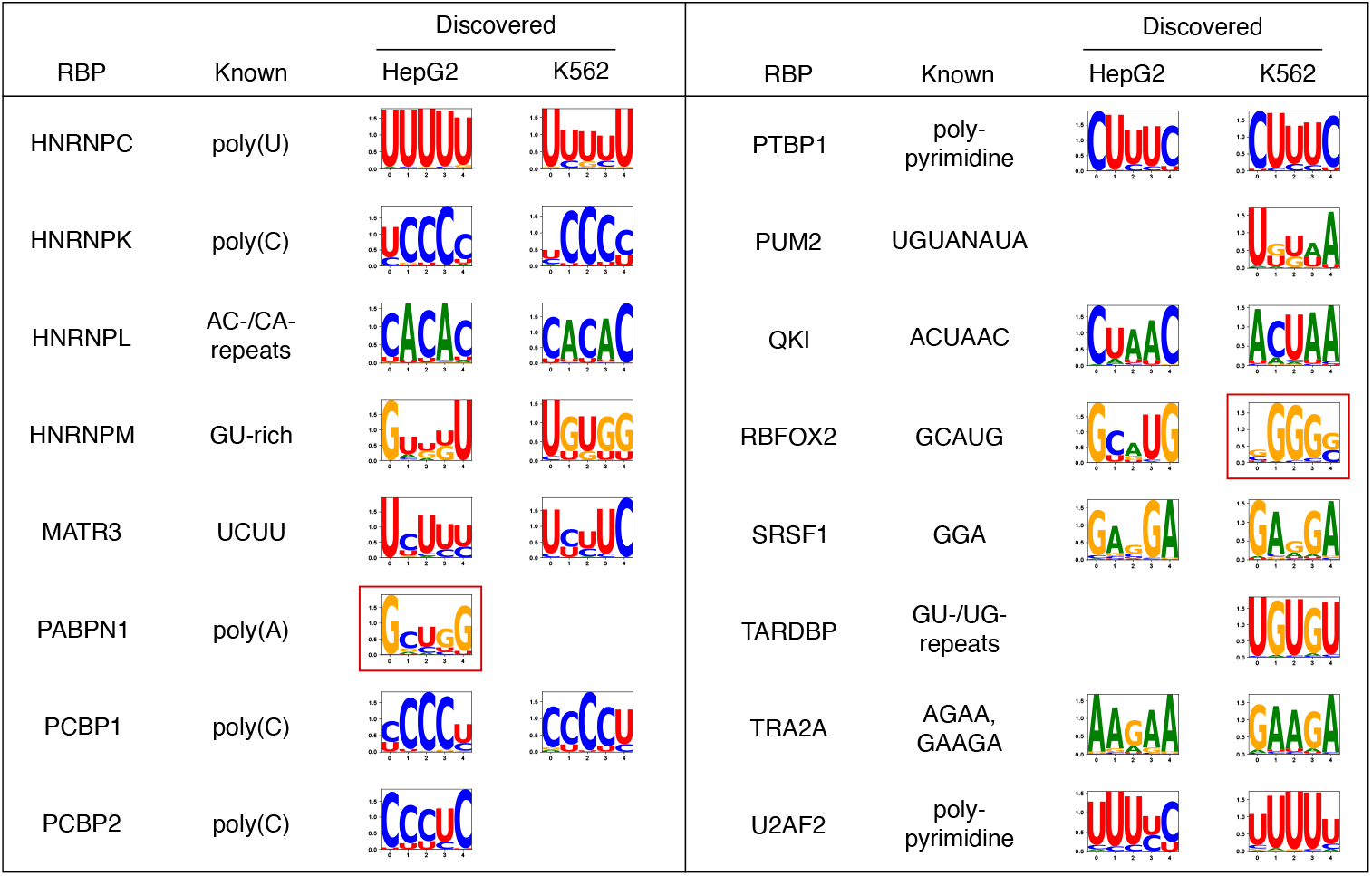
Discovered motifs for ground-truth set of RBPs in HepG2 and K562. For discovered motifs, only the primary motif selected by our iterative scoring strategy are displayed. Blank entries indicate that an eCLIP dataset was unavailable for the given RBP in the corresponding cell line (e.g., PUM2 in HepG2, PABPN1 in K562). Red boxes indicate discovered motifs that did not correspond with known motifs.

We achieved the same accuracy of 92.86% in K562; the consistency across cell lines demonstrates the robustness of our algorithm’s discovery. Our discovered motif for RBFOX2 in K562 was strongly G-rich. However, we note that our algorithm discovers all possible motifs in a given dataset, and thus we also discovered the canonical GCAUG motif despite it not being ranked as the primary motif (**Figure S1A**). This motif indeed resembles the known RBFOX2 motif and possesses similar features to the RBFOX2 motif discovered in HepG2; namely, a strong U preceding the canonical GCAUG 5-mer, and a strongly G-rich context (**Figure S1B, S1C**). In addition, we have previously demonstrated that our motif scoring method did not rank the canonical RBFOX2 motif as the primary motif in K562 due to the low enrichment of the consensus GCAUG, which helps to serve as an explanation for its selection of the more highly enriched G-rich motif as the primary motif [20].

### 3.2 Motif ranking demonstrates the importance of sequence context

As an additional form of validation, we applied STREME [21] to our datasets and compared the results against our discovered motifs (**Figure S2**). Our motif scoring strategy successfully selected the correct primary motif more often than did STREME’s ranking. In HepG2, our algorithm correctly identified the primary motif for 13 of 14 RBPs in the ground-truth set (with the exception being PABPN1), thus achieving 92.86% efficacy. By contrast, STREME correctly identified the primary motif for 11 of 14 RBPs, achieving 78.57% efficacy. First, STREME selected a G-rich motif as the primary motif for RBFOX2, ranking the canonical GCAUG motif in second place (**Supplemental Figure S3A**). Second, STREME selected a GCUGGAGUG motif as the primary for HNRNPC, and also ranked the canonical poly(U) motif in second place (**Supplemental Figure S4A**). The third misidentified primary motif belonged to PABPN1.

In K562, we observed similar outcomes (**Figure S2**). Our algorithm correctly selected the primary motif for 13 of 14 RBPs, with the exception of RBFOX2. However, STREME again failed to select the correct primary motif for both RBFOX2 and HNRNPC, again ranking first a G-rich motif and a GCUGGAGU motif, respectively, as it did in HepG2 (**Supplemental Figure S3B, S4B**).

We argue that *k*-mer enrichment in the context of RBP motifs is strongly related to these findings. First, RBFOX2 has been shown to bind in a G-rich sequence context *in vitro* [1] and *in vivo* [20]. We posit that the G-rich motif ranked first by STREME likely represents the sequence context of RBFOX2’s true motif GCAUG. Second, it is noteworthy that STREME selected the same primary motif, GCUGGAGU(G), for HNRNPC in both HepG2 and K562; the consistency in this finding across cell lines suggests that this motif may play a significant role in HNRNPC binding activity, either as an overrepresented *k*-mer in the context of HNRNPC’s poly(U) motif or as the motif for a different RBP that frequently interacts with HNRNPC. Our own motif discovery also supports this hypothesis: we discovered a GGAGU secondary motif in HepG2, as well as GAGUG and GGAGU secondary motifs in K562 (**Supplemental Figures S5, S6**). Taken together, these rankings demonstrate the significance and necessity of differentiating between motif and context *k*-mer units during the discovery process as well as their strong effect on the accuracy of motif discovery, as *k*-mers from flanking regions having high enrichment can be easily mistaken for candidate motif instances.

### 3.3 Our algorithm enables the discovery of RBP binding contexts and nucleotide preferences

We next generated our motif sequence contexts (**Figure 5**). Many of the discovered contexts and nucleotide preferences (**Figure 6**) correspond well with those from the literature: for example, RBFOX2 is known to bind in a G-rich environment; HNRNPK, PCBP1, and PCBP2 are known to bind in C-rich environments [22]; HNRNPC is known to bind poly(U) sequences [23]; and PTBP1 and U2AF2 are known to bind polypyrimidine tracts [24, 25]. Interestingly, we noted that the context nucleotide preferences for the same RBP across the two cell lines strongly resemble one another, indicating that, at least for these specific ground-truth sets, the RBPs have largely similar binding preferences.

**Fig. 5.**
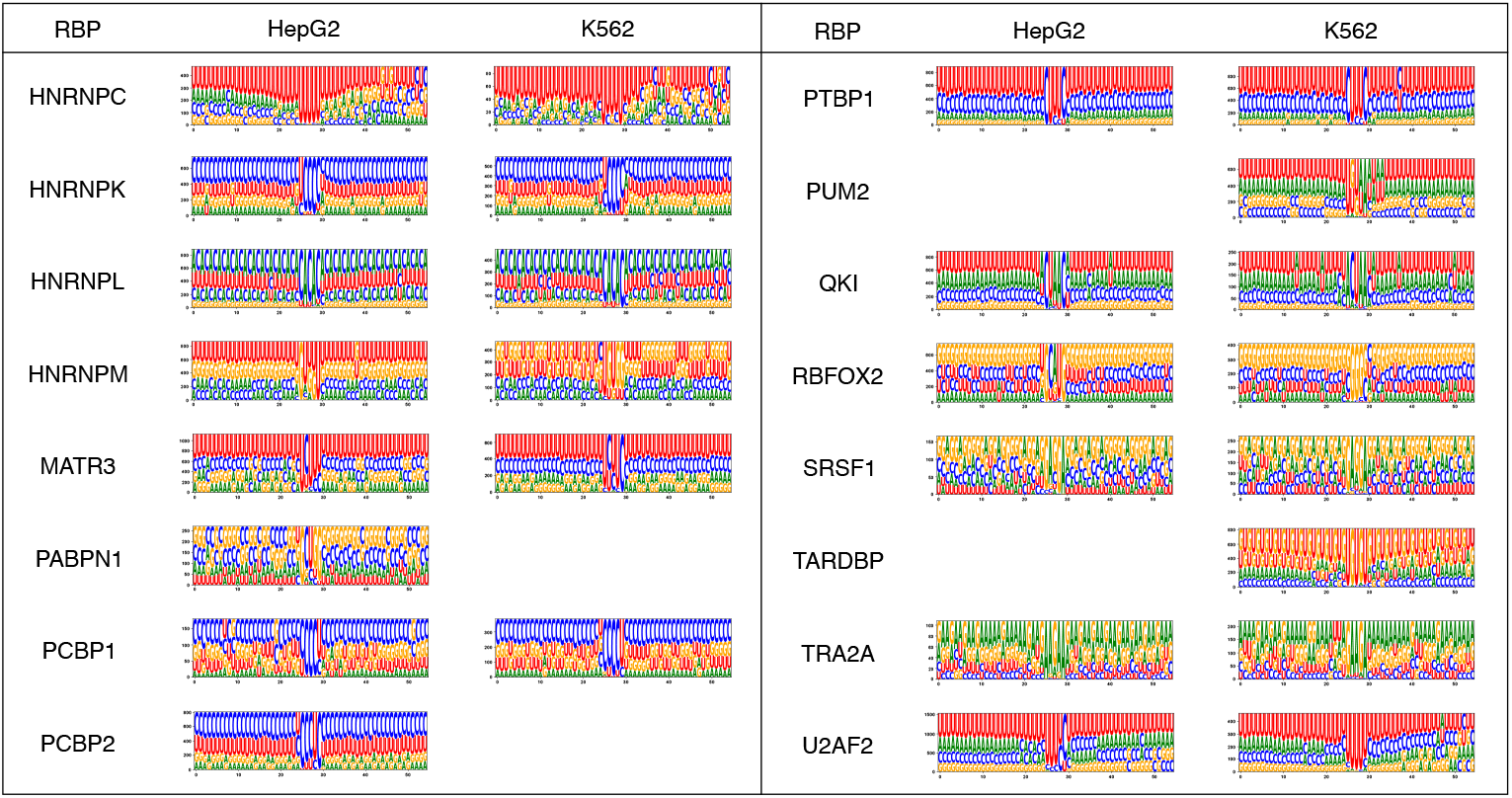
Discovered contexts of the primary motifs for the ground-truth set of RBPs in HepG2 and K562.

**Fig. 6.**
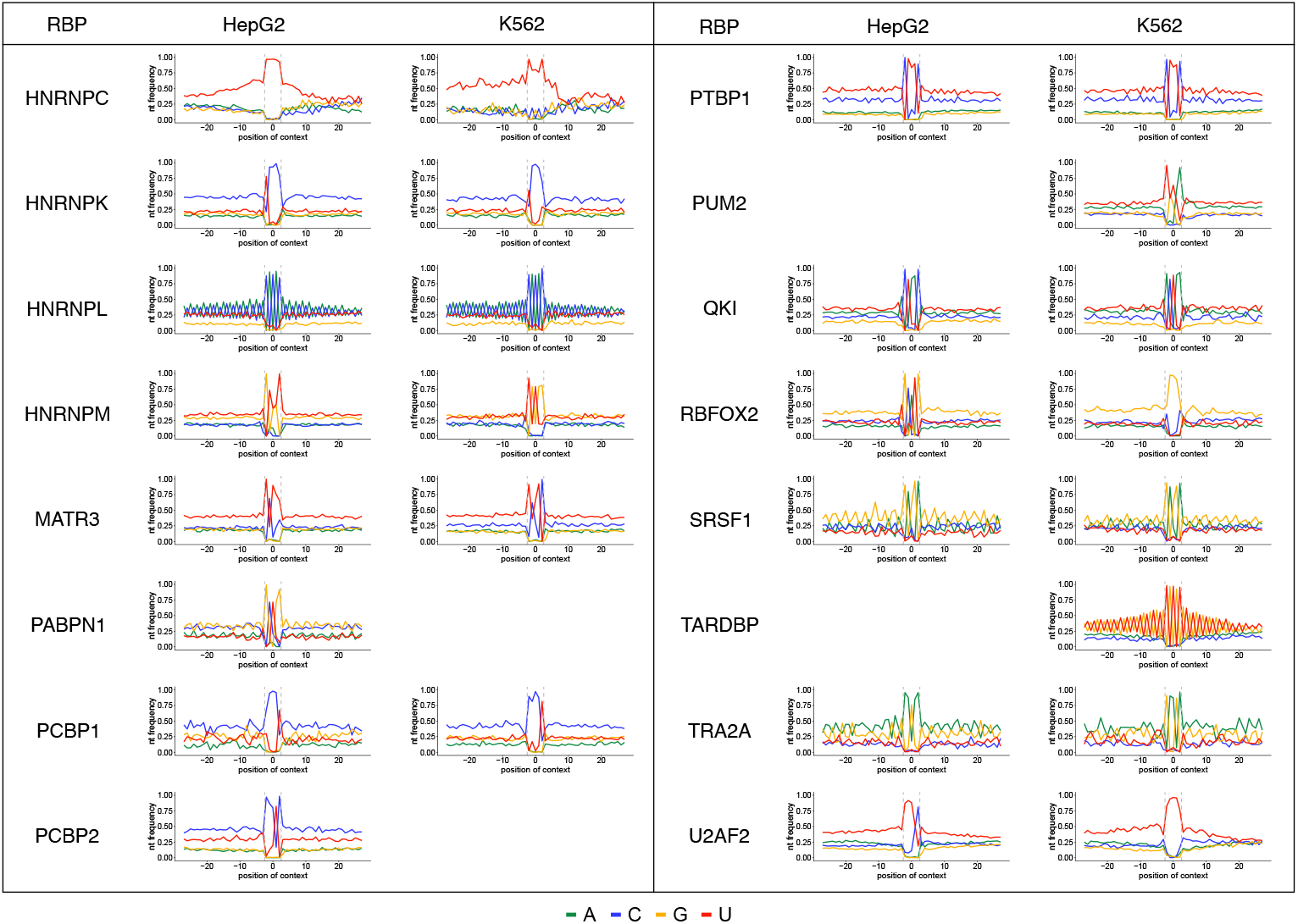
Discovered nucleotide preferences of the primary motifs for the ground-truth set of RBPs in HepG2 and K562.

To further investigate our hypothesis regarding HNRNPC secondary motifs, we examined the contexts of the secondary motifs. In both HepG2 and K562, when we inspected a longer context region of 101 nt, we observed that the most distal side of the left flanking region was strongly U-rich (**Supplemental Figures S5, S6**). This suggests that the GGAGU and GAGUG secondary motifs are located at the distal end of the right flanking region of HNRNPC’s primary poly(U) motif, and further supports our hypothesis that the secondary motifs represent contextual elements of HNRNPC’s canonical motif. Whether these secondary motifs are bound by other RBPs or *trans*-factors remains an open issue to be investigated.

The contexts logos and sequence preferences for the ground-truth sets presented herein serve as a preview for the potential, generalizability, and benefit of our algorithm. Indeed, we have applied our algorithm to large-scale eCLIP datasets across over 70 RBPs in each cell line, comprehensively characterizing the binding patterns of each [20]. In addition, we demonstrated the advantage of our method in discovering all possible motifs within a dataset, which allows investigation of potentially functional secondary motifs, the roles of motif clusters [20], and RBP-RBP interactions, thus aiding in the formation of new biological hypotheses regarding RBP-mediated RNA regulation.

## 4 Discussion

We have presented herein our consensus-based, context-aware, deterministic algorithm for the discovery of RBP motifs, contexts, and preferences. We described our linguistics-inspired formulation, which gives our model lexical, syntactic, and semantic layers, and specified the importance of *k*-mer enrichment, *k*-mer similarity, and *k*-mer co-occurrence as the three necessary and sufficient conditions for our algorithm’s accurate motif discovery. By validating our algorithm against established motifs from the literature for a set of well-characterized RBPs, we demonstrated the high accuracy of our algorithm. Furthermore, our algorithm not only demonstrated stability and consistency across the two cell lines, but it also enabled the discovery of novel RBP motifs, novel RBP sequence contexts, and the comprehensive characterization of binding preferences for a large array of diverse RBPs [20].

Our approach to motif discovery greatly reduced the search space at multiple stages. We achieved the first reduction in search space by using *k*-mer enrichments to compile a list of most highly-probable candidate consensuses, disregarding all other non-enriched *k*-mers in the motif discovery process. On average across all RBPs, this reduction led to only 33.8% of the 1024 possible *k*-mers being considered as candidate motif consensuses. We further reduced the search space during partition formation, in which we achieved an approximately 4.7-fold reduction by only considering *k*-mers with sufficient position-specific sequence similarity to the corresponding candidate consensus using our conditions, as opposed to a simple Hamming Distance requirement. In addition, we performed the co-occurrence tuning procedure for the top 10 most highly enriched candidate consensuses, which helped to reduce runtime.

We also introduced the consensus-instance co-occurrence as a novel and necessary attribute for motif discovery that may have sequence, structural, and biological interpretations. This property requires any given motif instance to occur within the same sequence as its corresponding motif consensus at some frequency; this frequency of co-occurrence is RBP-specific as well as consensus-specific, and it may reveal insights regarding the underlying sequence structure and biology of RBP binding. We posit potential biological and functional roles of the observed co-occurrence. Firstly, a high degree of co-occurrence between a motif consensus and a motif instance in the same sequence, especially when they have a larger Hamming distance, may signify a cooperative or competitive interaction between 2 RBPs. Secondly, a high degree of consensus-instance co-occurrence may also signify the possibility of homodimer formation and regulation by the RBP in question. This is particularly applicable to RBPs whose motif instances have a very small Hamming distance with the consensus. Finally, the co-occurrence relationship between motif consensus and motif instance is particularly intriguing in the disambiguation of genomic language, as word co-occurrence is an important linguistic concept that provides a measure of word association and semantic relation between two lexical units [8, 9]. Future work is needed to improve our understanding of consensus-instance co-occurrence.

Our algorithm’s implementation is easily parallelizable, which greatly improves runtime and efficiency. Another advantage of our algorithm is its discovery of all possible motifs within a given dataset, which means that even if our scoring method does not rank the true motif as the primary, the algorithm is still able to detect the true motif, as we demonstrated with examples above. This will enable investigation of secondary motifs that may have important, specialized biological roles. Additionally, because we are able to identify each distinct motif instance, it would be interesting to examine whether the sequence contexts and their nucleotide preferences vary for specific motif instances that comprise the motif. Finally, we foresee multiple avenues in which the algorithm can be applied to the context data structure without having to rely on MIL predictions, which would further increase its versatility.

## Data availability

Data used in this work originate from ENCODE eCLIP merged replicate peak files, which are available at the ENCODE portal [26–28].

## Code Availability

Code is publicly available at https://github.com/orbitalse/novel RBP discovery methods.

## Author Contributions

S.I.E.: Problem Formulation, Conceptualization, Data Curation, Data Processing, Investigation, Formal Analysis, Methodology, Visualization, Writing of Manuscript (original draft), Review of Manuscript, Editing of Manuscript. Z.W.: Research Area, Discussions, Review of Manuscript. All authors reviewed and approved the final version of the manuscript.

**Supplementary Figure S1.**
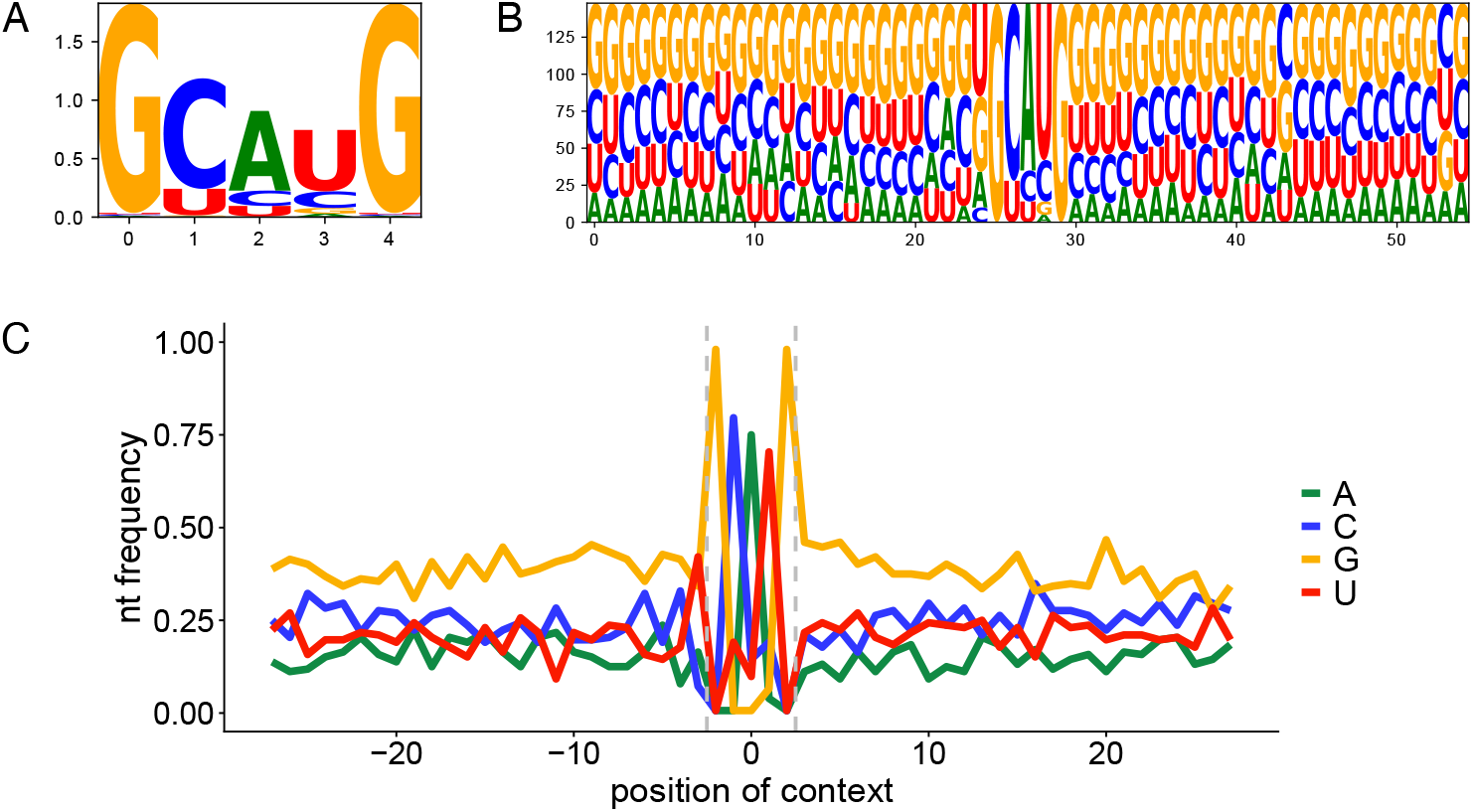
Motif, context, and nucleotide preference discovery for the GCAUG consensus for RBFOX2 in K562. (A) Discovered GCAUG motif corresponds with the canonical RBFOX2 motif and resembles the GCAUG motif discovered in HepG2. (B) Discovered context logo, including a U preceding the GCAUG consensus. (C) Nucleotide preferences for the GCAUG motif, demonstrating a G-rich preference.

**Supplementary Figure S2.**
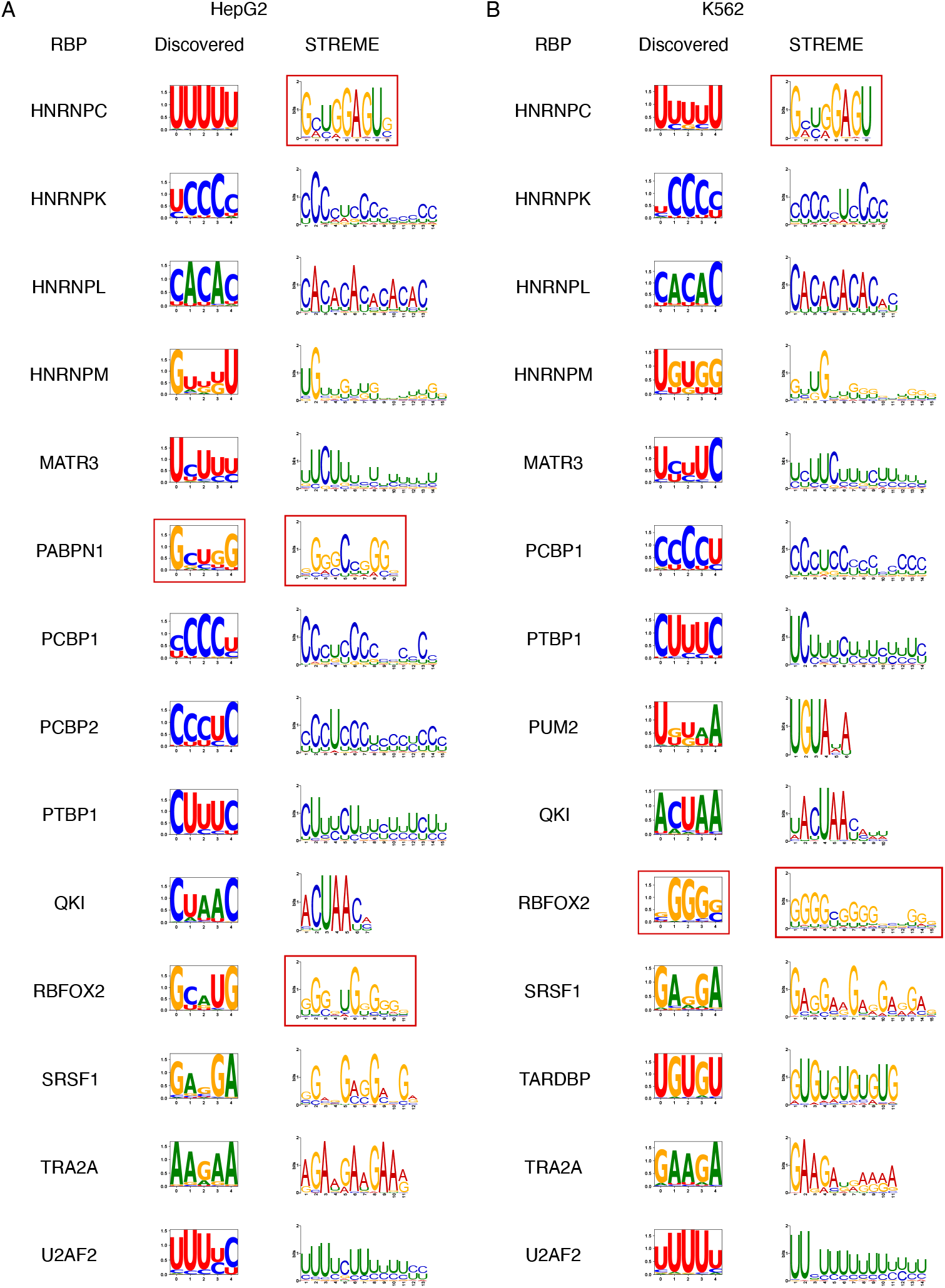
Comparison of primary motifs discovered by our algorithm and by STREME for the ground-truth sets in (A) HepG2 and (B) K562. Misidentified primary motifs by each method are outlined in red.

**Supplementary Figure S3.**
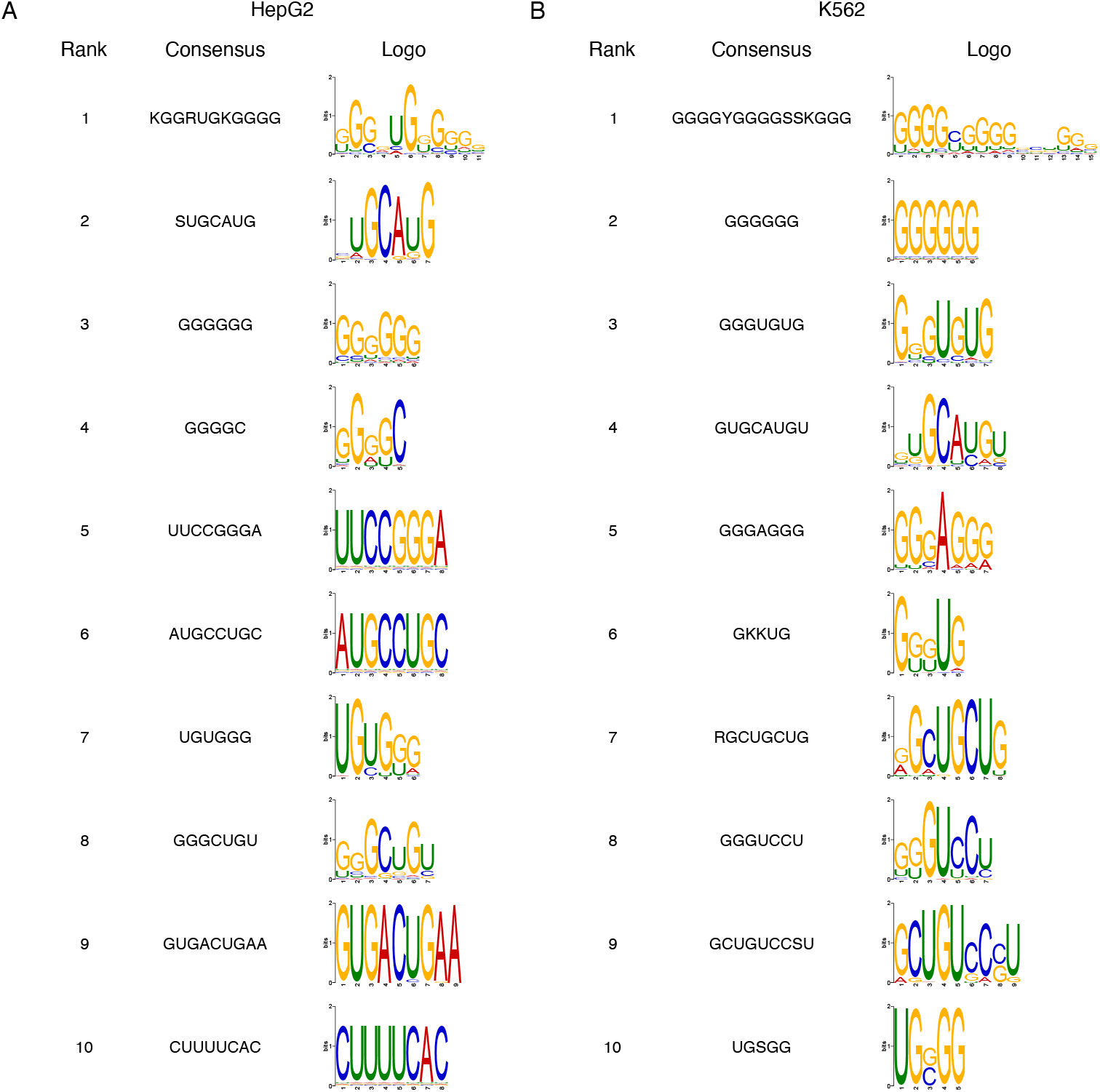
Ranked motifs discovered by STREME for RBFOX2 in (A) HepG2 and (B) K562.

**Supplementary Figure S4.**
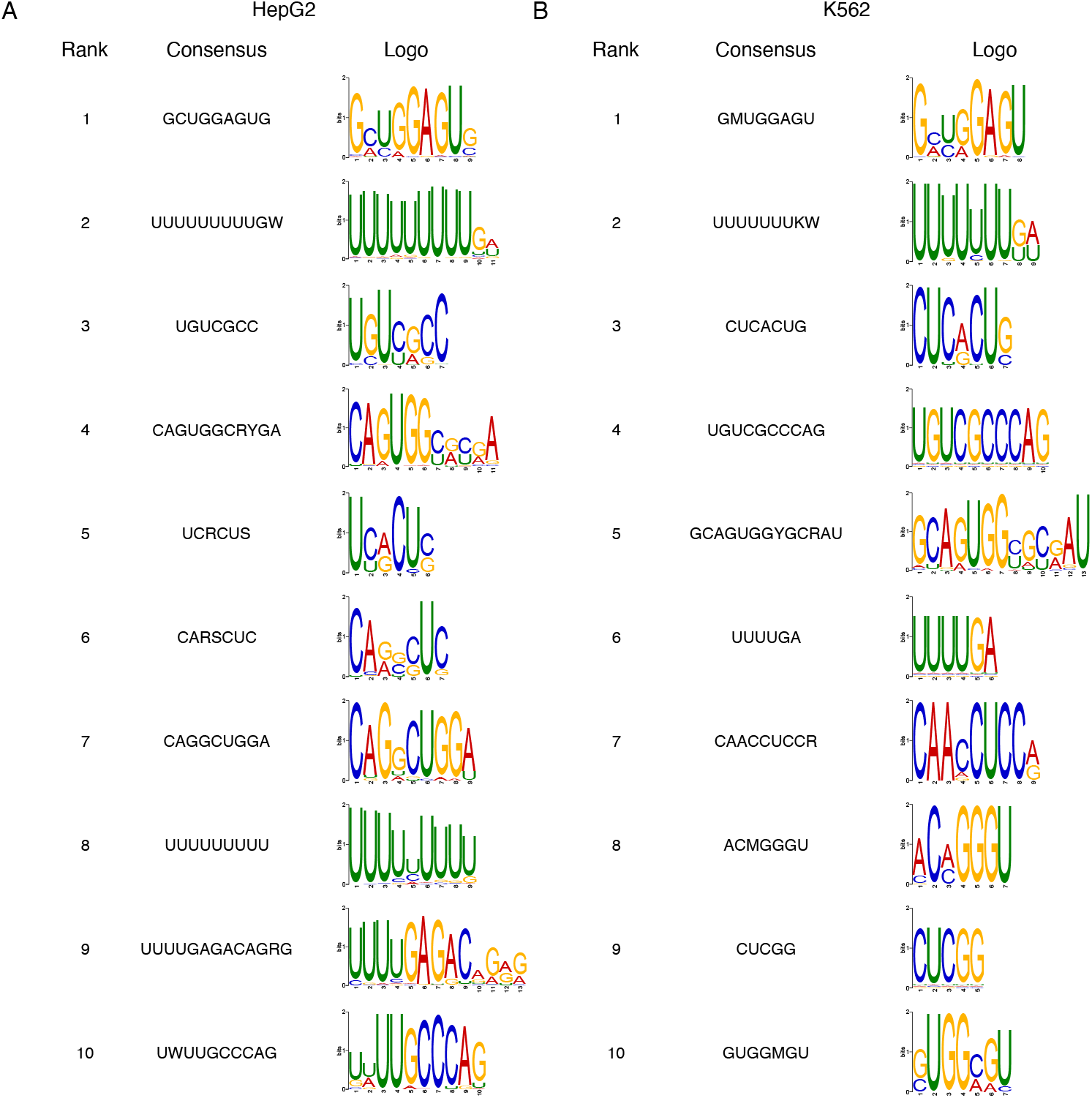
Ranked motifs discovered by STREME for HNRNPC in (A) HepG2 and (B) K562.

**Supplementary Figure S5.**
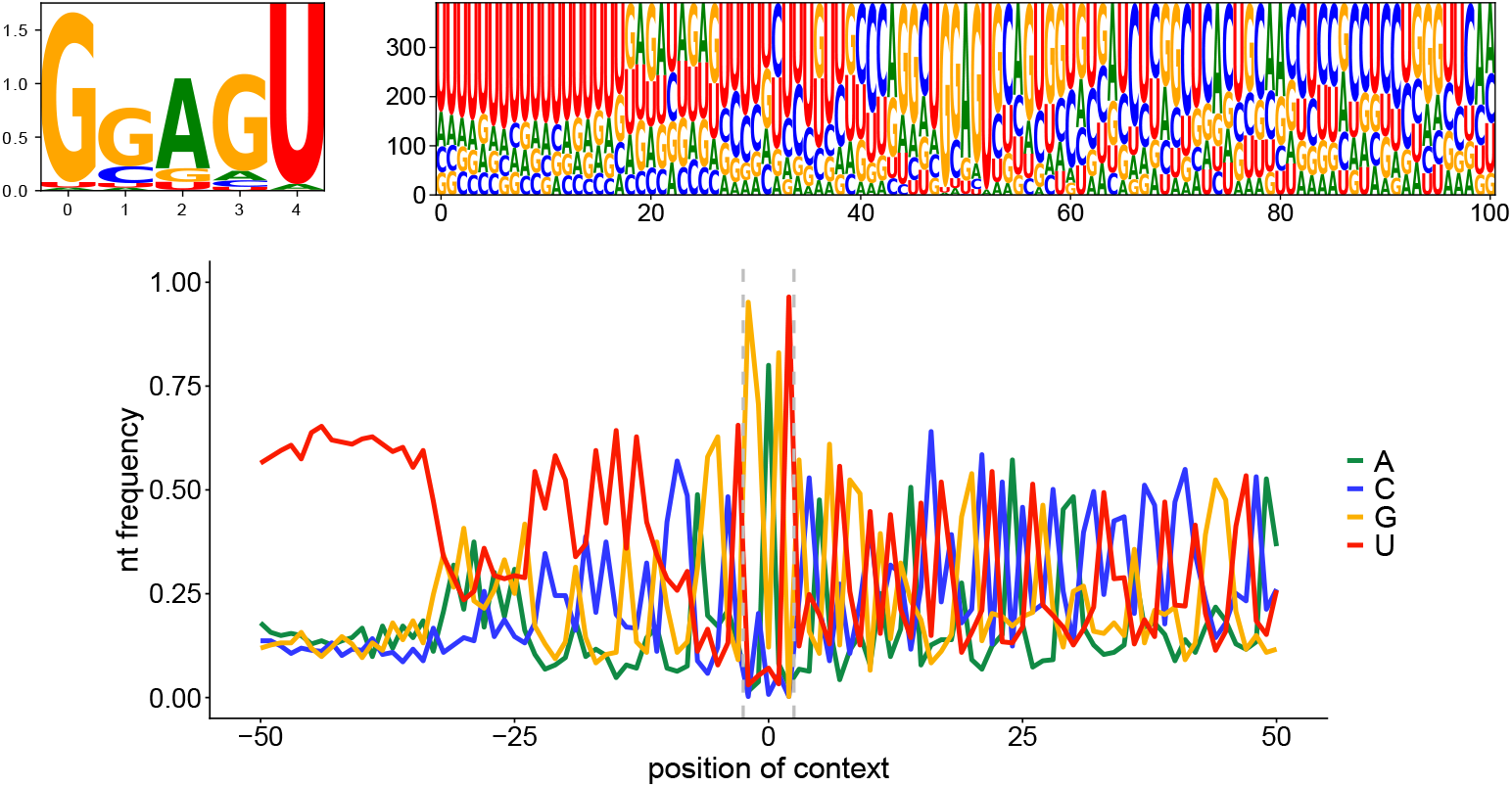
Secondary GGAGU motif for HNRNPC in HepG2, along with its context logo and nucleotide preferences.

**Supplementary Figure S6.**
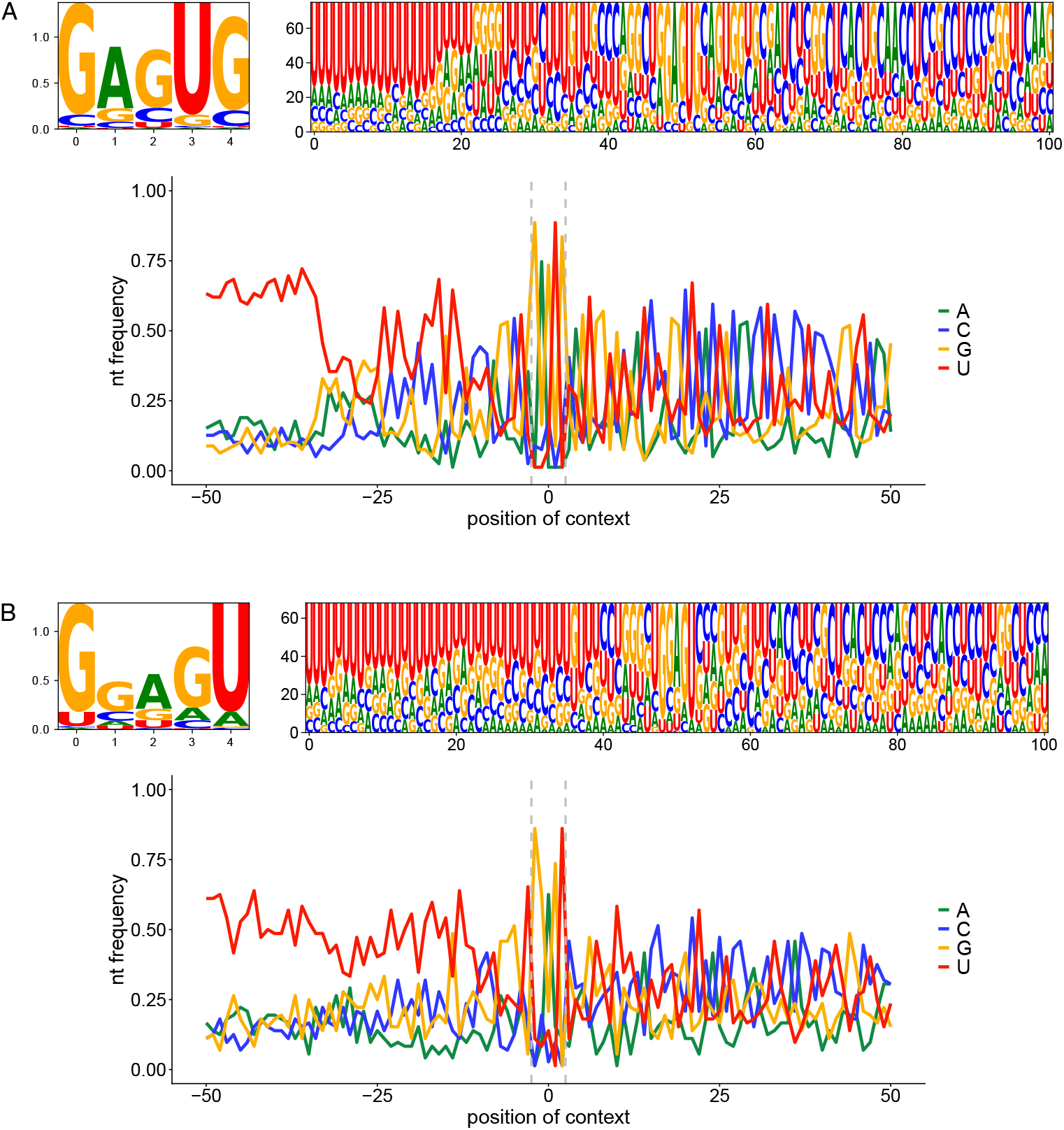
Secondary motifs for HNRNPC in K562, along with their respective context logos and nucleotide preferences. (A) Secondary GAGUG motif. (B) Secondary GGAGU motif.

### Algorithm 1

Motif Discovery Algorithm

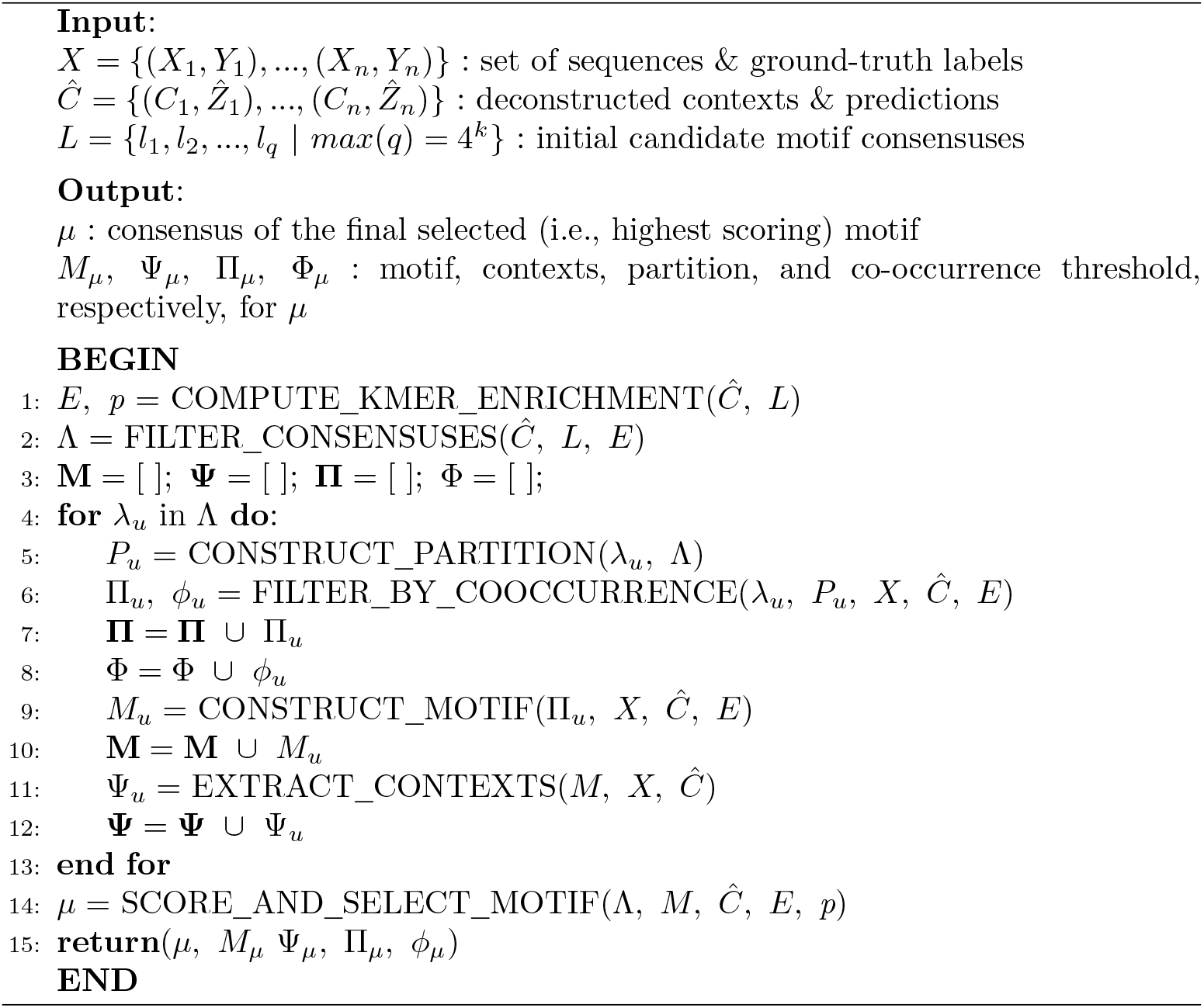

### Algorithm 2

Co-occurrence Filtering

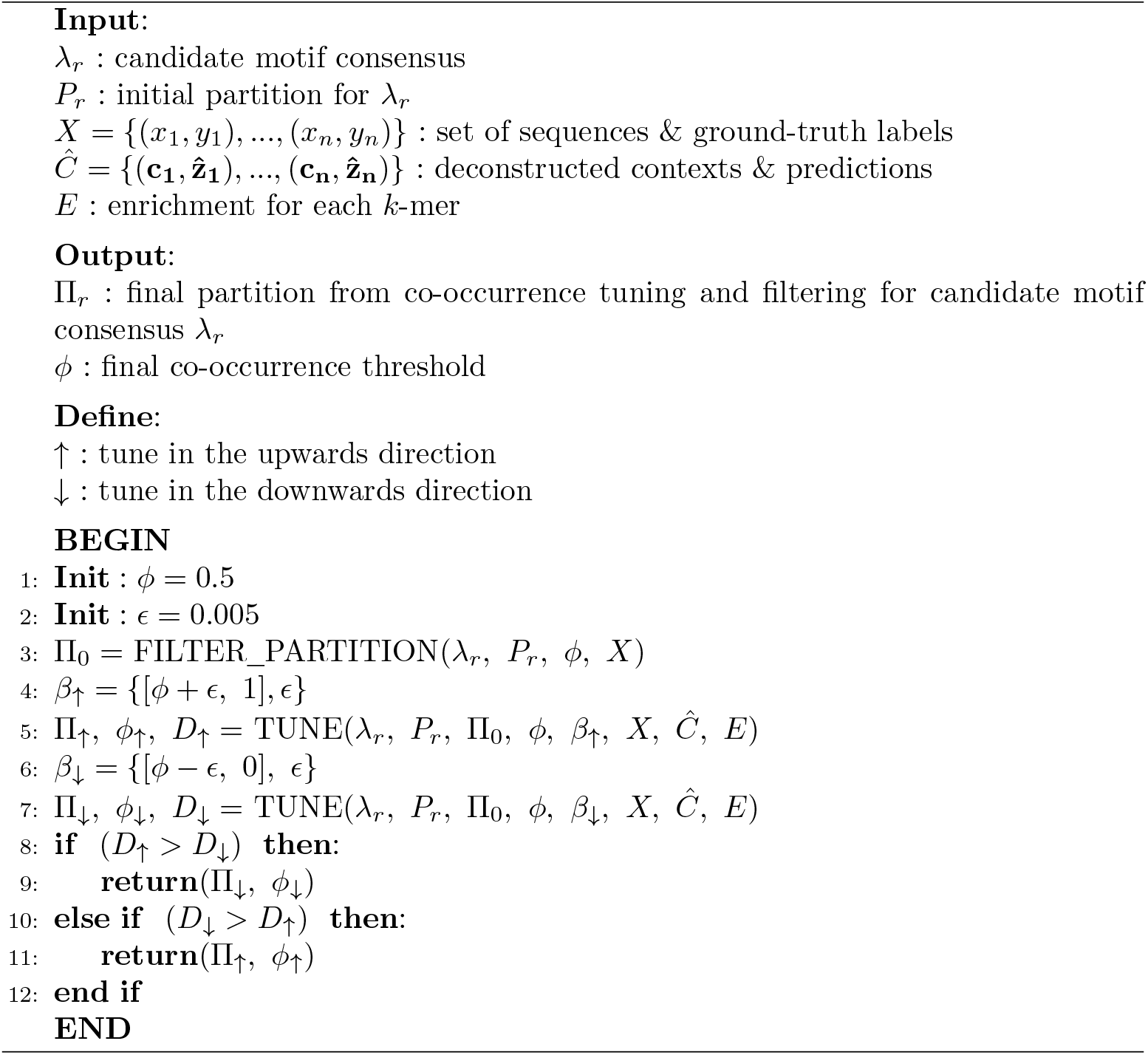

### Algorithm 3

Co-occurrence Tuning

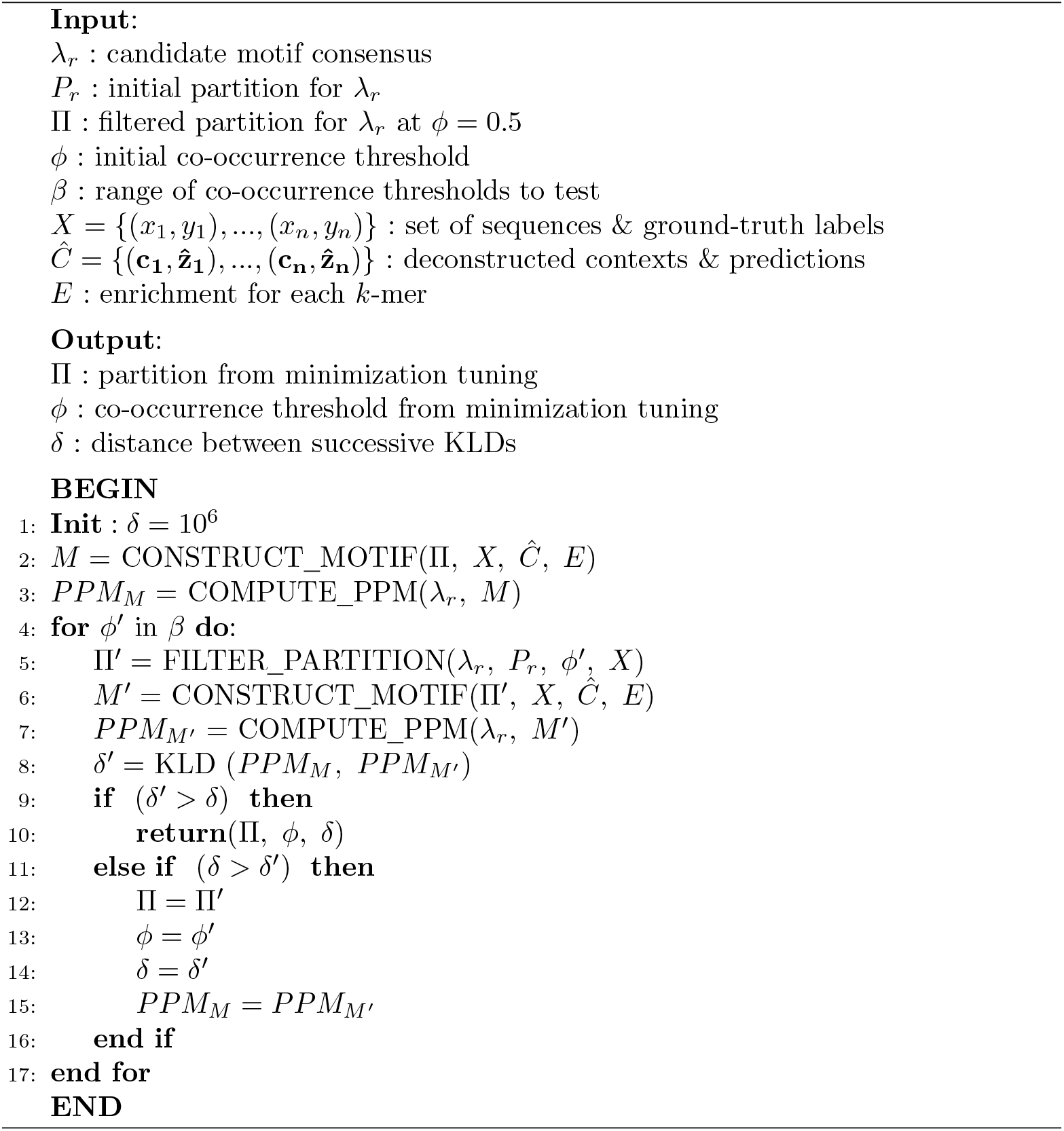

### Algorithm 4

Primary Motif Selection by Multi-metric, Iterative Scoring

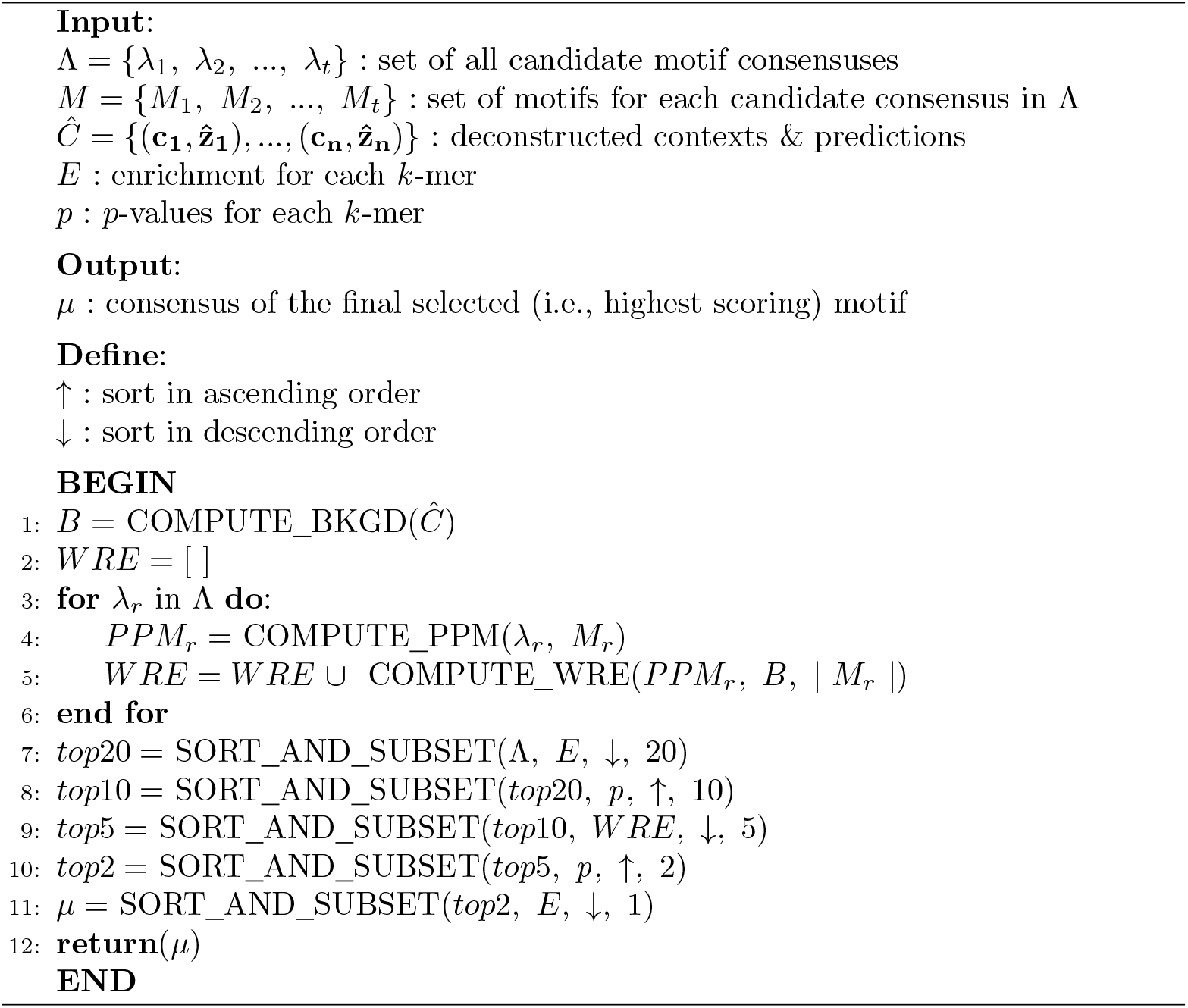

